# Integrated computational and *Drosophila* cancer model platform captures previously unappreciated chemicals perturbing a kinase network

**DOI:** 10.1101/344192

**Authors:** Peter Man-Un Ung, Masahiro Sonoshita, Alex P. Scopton, Arvin C. Dar, Ross L. Cagan, Avner Schlessinger

**Author notes:** These authors contributed equally to this work (Peter Man-Un Ung, Masahiro Sonoshita).

## Abstract

*Drosophila* provides an inexpensive and quantitative platform for measuring whole animal drug response. A complementary approach is virtual screening, where chemical libraries can be efficiently screened against protein target(s). Here, we present a unique discovery platform integrating structure-based modeling with *Drosophila* biology and organic synthesis. We demonstrate this platform by developing chemicals targeting a *Drosophila* model of Medullary Thyroid Cancer (MTC) with disease-promoting kinase network activated by mutant dRet^M955T^. Structural models for kinases relevant to MTC were generated for virtually screening to identify initial hits that were dissimilar to known kinase inhibitors yet suppressed dRet^M955T^-induced oncogenicity. We then combined features from the hits and known inhibitors to create a ‘hybrid’ molecule with improved dRet^M955T^ phenotypic outcome. Our platform provides a framework to efficiently explore novel chemical spaces, develop compounds outside of the current inhibitor chemical space, and ‘correct’ cancer-causing signaling networks to improve disease prognosis while minimizing whole body toxicity.

**AUTHOR SUMMARY:** Effective and safe treatment of multigenic diseases often involves drugs that modulate whole systems by interacting with multiple nodes in pathways and networks, i.e., polypharmacology. Polypharmacology is increasingly appreciated as a potential desirable property of kinase drugs; however, most known drugs that interact with multiple targets have been identified as such by chance, and most polypharmacological compounds are not chemically unique resembling to structures of known kinase inhibitors. The fruit fly *Drosophila* has been established as a robust screening platform that provides an inexpensive, rapid, and quantitative measure of whole animal drug response, complementing computational approaches. We present a chemical genetics approach that efficiently combines *Drosophila* with structural prediction and virtual screening, creating a unique discovery platform. We demonstrate the utility of our approach by developing useful small molecules targeting a kinase network in a *Drosophila* model of Medullary Thyroid Cancer (MTC) driven by the active mutant dRet^M955T^.

## INTRODUCTION

Protein kinases play a key role in cell signaling and disease networks and make up major therapeutic targets. The limited capacity to test large number of compounds exploring diverse chemical scaffolds, coupled with the low translatability of *in vitro* kinase inhibition into whole animal efficacy, effectively constrain the chemical space of the known kinase inhibitors (KIs). Thus, obtaining optimal KIs at clinically relevant therapeutic levels is challenging, despite extensive academic and industry effort.

To expand the number of kinase inhibitors, a variety of platforms have recently emerged as useful tools for compound screening. *Drosophila melanogaster* (fruit fly) provides an inexpensive and efficient biological platform for cancer drug screening, capturing clinically relevant compounds [1–3]. For example, *Drosophila* was used to help validate vandetanib as a useful treatment for medullary thyroid cancer [4] (MTC). As a screening platform, *Drosophila* offers several advantages: First, flies and humans share similar kinome and kinase-driven signaling pathways [5], facilitating the use of flies to predict drug response in humans [1, 6]. Second, the ease of breeding and the short (~10 day) life cycle of *Drosophila* make it possible to carry out efficient mid-throughput chemical screening in a biological system. Third, the screening readout provides a quantitative animal-based measurement of structure-activity relationships (SAR), and further provides information on the therapeutic potential or toxicity of the tested compounds: measurable parameters include survival and multiple phenotypic indicators that depend on kinase activity.

One key limitation of *Drosophila-based* mid-throughput screening platform is that it cannot explore very large chemical libraries [7], such as the ZINC library which has over 750 million purchasable compounds [8]. In contrast, structure-based virtual screening is a fast and inexpensive computational method that can screen large compound libraries, a useful approach to identify unique chemical probes [9]. If the structure of the protein is unknown, virtual screening can be performed against the homology models of the target constructed based on experimentally determined structures. However, the automated construction of homology models—with sufficient accuracy for virtual screening for multiple targets simultaneously and the application of molecular docking to signaling networks—remains challenging in particular for highly dynamic targets such as kinases [10, 11] and would benefit from a readily accessible whole animal platform.

RET is a receptor tyrosine kinase associated with multiple roles in development and homeostasis. Activation of RET by the mutation M918T (analogous to *Drosophila* M955T) is associated with MTC [12, 13]. Transgenic *Drosophila* expressing the dRet^M955T^ isoform show key aspects of transformation, including proliferation and some aspects of metastasis [6, 14]. Genetic modifier screens with dRet^M955T^ flies led to the identification of multiple RET pathway genetic ‘suppressors’ and ‘enhancers’, loci that when reduced in activity improve or worsen the disease phenotype, respectively. These functional mediators of RET-dependent transformation include members of the Ras/ERK and PI3K pathways as well as regulators of metastasis such as SRC [6, 15].

Oral administration of the FDA approved, structurally related multi-kinase inhibitor analogs sorafenib and regorafenib, along with additional structural analogs, partially rescued *dRet*^*M955T*^*-induced* oncogenicity in *Drosophila* [1, 15]. Sorafenib class inhibitors are classified as ‘type-II’ KIs that bind the kinase domain in its inactive state [16], a conformational state regulated by the aspartate-phenylalanine-glycine (DFG)-motif (Fig. 1A) [17, 18 17, 18 17, 18 17, 18 17, 18 17, 18]. In the inactive, ‘DFG-out’ conformational state the directions of DFG-Asp and DFG-Phe ‘flip’, vacating a pocket previously occupied by DFG-Phe (‘DFG-pocket’) that modulates binding to type-II inhibitors. A key challenge of targeting kinases in the DFG-out conformation with structure-based virtual screening is that few kinase structures have been reported with the DFG-out conformation [19]. We have recently developed DFG-model [10], a computational method for modeling kinases in DFG-out conformations. This method informed the mechanism of clinically relevant multi-kinase inhibitors that target the MTC network [15].

**Figure 1.**
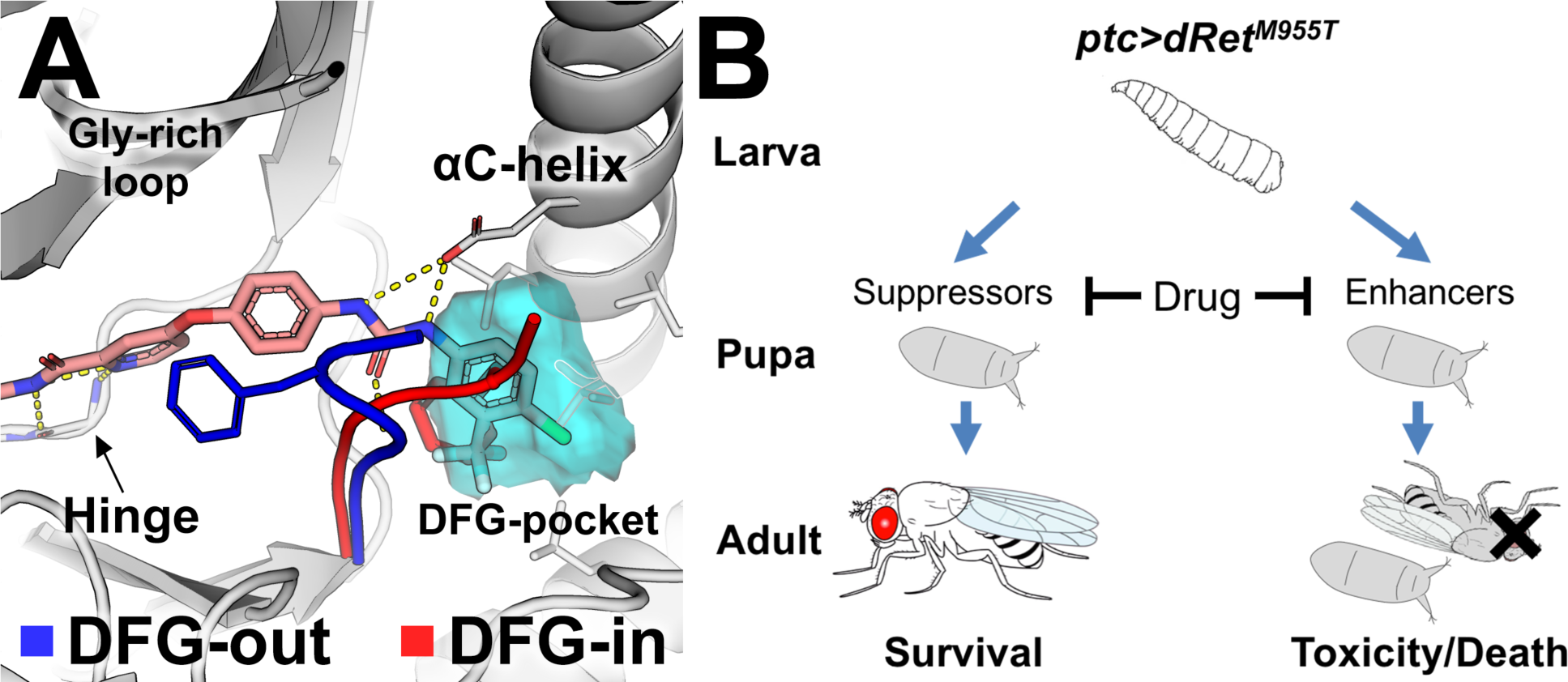
Kinase binding to type-II kinase inhibitors. ***(A)*** The conformational state of protein kinases (e.g., KDR) including DFG-in (red) and DFG-out (blue) is determined by the DFG-motif. The DFG-pocket (cyan mesh) is unique to the DFG-out conformation. Sorafenib is shown in pink. Broken yellow lines indicate hydrogen bonds. ***(B)*** A scheme depicting the positive and negative effects of drug acting on genetic modifiers of medullary thyroid cancer in a *Drosophila* model. ptc-driven *dRet*^*M955T*^ induces lethality during development. ‘Suppressors’ or ‘enhancers’ suppress or enhance, respectively, *dRet*^*M955T*^*-in*duced disease phenotypes as revealed in genetic screening. A drug can suppress lethality by inhibiting the suppressors. It can also induce toxicity and/or worsen transformed phenotypes by inhibiting the enhancers, which results in enhanced lethality.

In this study, we report the development of an integrated platform (Fig. 2) that combines (i) computational modeling of kinases in their inactive state plus massive multitarget virtual screenings with (ii) whole animal *Drosophila* assays to discover previously unappreciated chemicals that perturb the RET-dependent transformation. Furthermore, we leverage these insights to create a novel ‘hybrid’ molecule with unique chemical structure and biological efficacy. Finally, we discuss the relevance of this approach to expedite the discovery of novel chemical scaffolds targeting disease networks.

**Figure 2.**
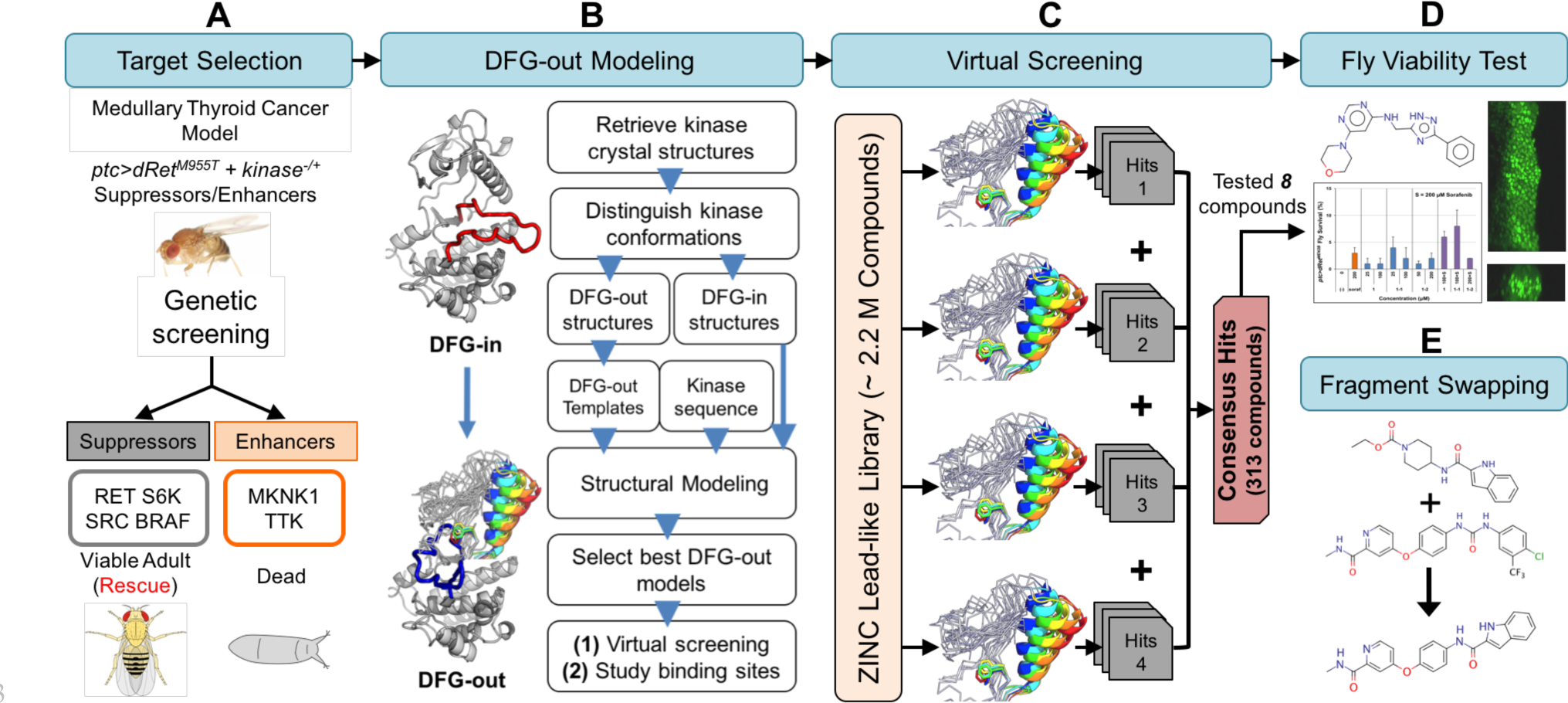
Fly genetics and computational chemistry discovery platform. Key steps include **(A)** determining suppressors and enhancers in a dominant modifier genetic screening and their *in silico* modelability, **(B)** generating DFG-out kinase models using DFGmodel, **(C)** virtual screenings of compound libraries against the modeled suppressors and enhancers, combining top-ranking screening results into consensus result, **(D)** testing top-ranking compounds for rescue of lethality (left panel) and migration of transformed cells in developing wing discs of *ptc>dRet*^*M955T*^ flies (right panel), and **(E)** refining hits by combining structural elements of computationally derived hits and that of drugs and evaluating new targets.

## RESULTS

### Target selection from fly genetic screen and structural analysis

In transgenic *patched-GAL4;UAS-dRet^M955T^ (ptc>dRet^M955T^)* flies, the *ptc* promoter drives expression of an oncogenic isoform of *Drosophila* Ret in multiple tissues; the result is lethality prior to adult eclosion [1, 15]. We previously used this and similar fly MTC models in genetic screens to identify 104 kinases that mediate *dRet*^*M955T*^*-mediated* transformation [15] (Figs. 1B, 2A, S1).

To narrow this list, we prioritized candidates based on two considerations: (i) pharmacological relevance - kinases downstream of RET signaling were prioritized due to their established functional role [6, 20]; (ii) structural coverage - kinases with known DFG-out structures or those that can be modeled with sufficient accuracy in this conformation were further investigated [10]. Atypical kinases (*e.g.*, mTOR and eEF2K) and members of the RGC family were excluded as they have diverse sequence and structure features that limit our ability to generate accurate homology models. Applying these criteria to our genetic modifier list, we focused on targeting four key kinase targets: RET (receptor tyrosine kinase), SRC (tyrosine kinase), BRAF (tyrosine kinase-like), and p70-S6K (AGC family).

### Modeling kinases in DFG-out conformation

Description of the various conformations adopted by the kinases during activation and inhibition is needed for rationally designing novel, conformation-specific inhibitors. Therefore, our approach was to perform massive structure-based virtual screenings of purchasable compound libraries against multiple models with DFG-out conformation; our goal was to identify generic kinase inhibitors that may target one or more prioritized kinases - but more importantly - have an effect on the disease pathway in the animal model.

The structure of two of the kinases identified in our *dRet*^*M955T*^ model—BRAF and SRC—have been solved in the DFG-out conformation; the DFG-out structures of RET and p70-S6K have not been reported. We therefore generated DFG-out models using DFGmodel, a computational tool that generates homology models of kinase in DFG-out conformation through multiple-template modeling that samples a range of relevant conformations [10]. These models enrich known type-II inhibitors among a diverse set of non-type-II KIs found in PDB with accuracy similar to or better than that obtained for experimentally determined structures and provides approximation for binding site flexibility [10]. For example, in a recent application of DFGmodel, models generated by this method were used in parallel with medicinal chemistry to optimize clinically relevant compounds that are based on the established kinase inhibitor sorafenib [15]. Conversely, in this study, models generated by DFGmodel are used to develop compounds that are outside of the current kinase inhibitor chemical space.

To guide the identification of a “generic” kinase inhibitor of a disease pathway, we first compared the DFG-out models of the kinases, identifying key similarities and differences in physicochemical properties among their inhibitor-binding sites. First, we noted that the prioritized targets RET, BRAF, p70-S6K, and SRC present negative electrostatic potential on the DFG-pocket surface, while many non-targets such as ERK have positive electrostatic potential (Fig. 3A). This difference may partially explain the partial selectivity of type-II inhibitors (*e.g.*, sorafenib) toward our prioritized targets but not on electrostatic positive kinases such as ERK. Second, RET and SRC have large DFG-pocket volumes (163 Å^3^, 196 Å^3^); p70-S6K and BRAF have moderately large pockets (158 Å^3^, 136 Å^3^). In contrast, ERK has a small DFG-pocket (113 Å^3^) (Fig. 3B). We used this size difference to computationally select for kinases with larger DFG-pockets (e.g., RET, SRC) while excluding kinases with smaller DFG-pockets (*e.g.*, ERK).

**Figure 3.**
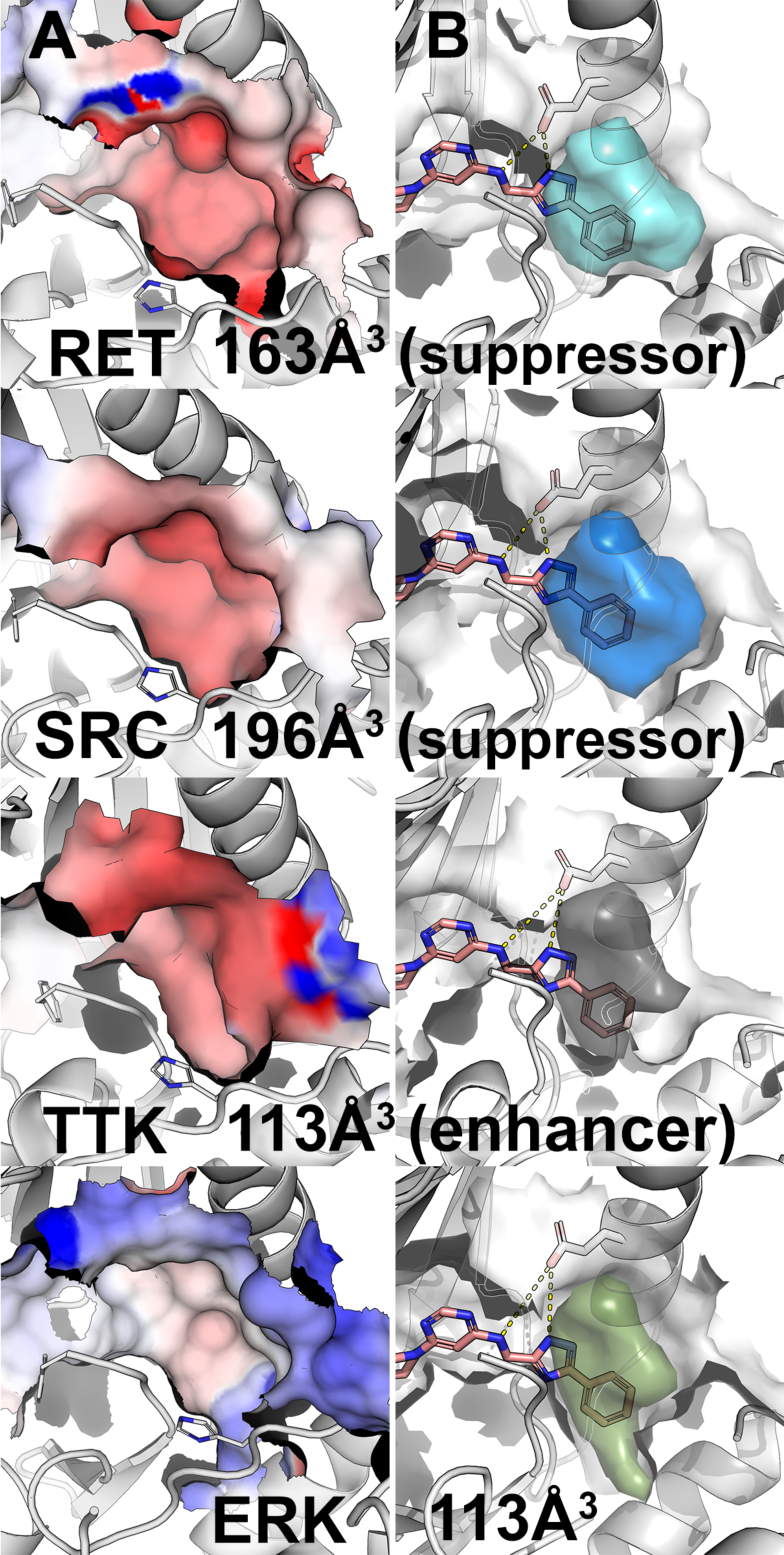
Visualization of DFG-pockets. **(A)** Electrostatic potential (red, negative potential; blue, positive potential) on the surface of the DFG-pocket in various kinases, including the suppressors RET and SRC, the enhancer TTK, and ERK. **(B)** Accessible volume of the DFG-pocket (colored volume) for potential type-II kinase inhibitor. Hit molecule *1* is depicted in pink sticks. Broken yellow lines indicate hydrogen bonds.

### Virtual screening against MTC pathway

We performed virtual screening against multiple DFG-out models of MTC targets to identify putative small molecules that modulate the disease network (Fig. 2C). We docked a purchasable lead-like library from the ZINC database [21] (2.2 millions compounds) against 10 DFG-out models for each kinase target, yielding over 88 million total docking poses. To combine the screening results, a two-step consensus approach was used. In the first step, compounds that ranked in the top 10% in 5 or more of the 10 models of each kinase were selected, resulting in approximately 2,000 compounds per kinase. In the second step, compounds that ranked in the top 25% in at least 3 of 4 targets were selected, resulting in 247 compounds. For comparison, sorafenib, an inhibitor that rescues *ptc>dRet*^*M955T*^ flies, would rank eighth in this consensus docking result. From these consensus compounds, eight commercially available compounds were purchased to test their ability to rescue *ptc>dRet*^*M955T*^ flies (Table S1). These compounds were selected based on their interactions with key elements of the “ensemble” of targets’ binding sites, with the emphasis on the conserved glutamate in αC-helix, the amide backbone of DFG-aspartate, and if present, the amide backbone of the hinge region (Fig. S3). Although the compounds are not predicted to bind optimally to each one of our targets, we hypothesized that these compounds may have an additive effect on the disease pathway, which could be improved with medicinal chemistry.

### Testing candidates in ptc>dRet^M955T^ fly viability assay

Transgenic *ptc>dRet*^*M955T*^ flies express the oncogenic *Drosophila* dRet^M955T^ isoform in several tissues in the developing fly, leading to transformation of dRet^M955T^ tissues [6, 14]. As a result, *ptc>dRet*^*M955T*^ flies exhibited 0% adult viability when cultured at 25°C, providing a quantitative ‘rescue-from-lethality’ assay to test drug efficacy [1, 15]. Compounds were fed at the highest accessible concentrations (see Experimental Procedures). We used sorafenib as a positive control, as it previously demonstrated the highest level of rescue among FDA-approved KIs in *ptc>dRet*^*M955T*^ flies [15]. Similar to our previous results, feeding *ptc>dRet*^*M955T*^ larvae sorafenib (200 μM) improved overall viability to 3-4% adult survival (*P* < 0.05).

We used this rescue-from-lethality assay to test the efficacy of the eight compounds identified through virtual screening (Figs. 4B, 5B). When fed orally, two unique compounds, ***1*** and ***2*** (Table S2), rescued a small fraction of *ptc>dRet*^*M955T*^ flies to adulthood (Figs. 4A and 5A) and did not affect the body size of the larvae and pupae, a metrics for comparing food intake, of *ptc>dRet*^*M955T*^ flies when compared to the wild-type. At the maximum final concentration in fly food (100 μM), ***1*** rescued 1% (P < 0.05) *ptc>dRet*^*M955T*^ flies to adulthood as compared to 3-4% rescue by sorafenib at 200 μM (Fig. 4A). ***1*** is characterized by the 3-phenyl-(1H)-1,2,4-trazole moiety (Fig. 4B). **2**, characterized by the 1 H-indole-2-carboxamide moiety, improved *ptc>dRet*^*M955T*^ fly viability to an average of 1 % (P < 0.05) when tested at 25-400 μM (Fig. 5A, B).

**Figure 4.**
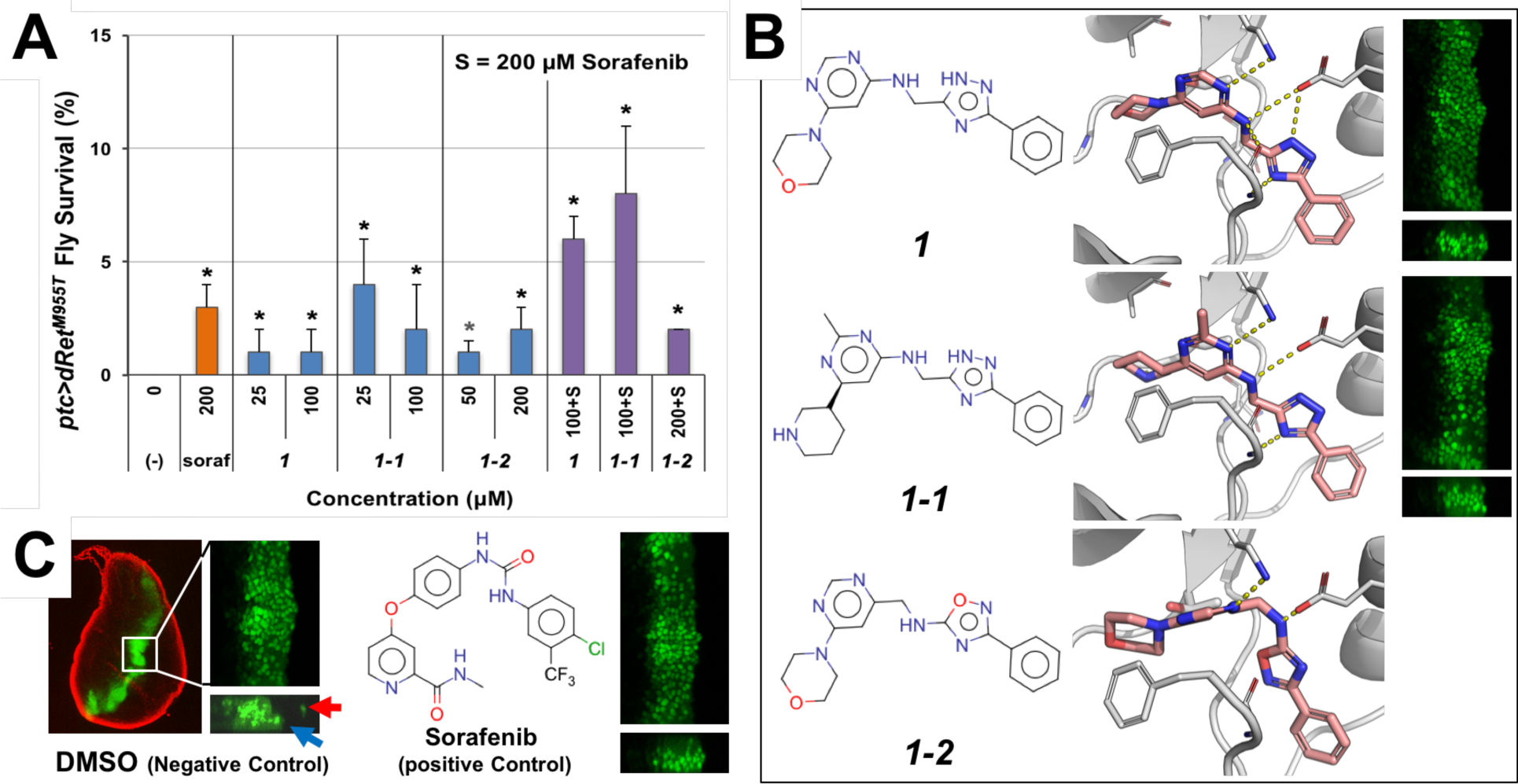
Compound *1* and its analogs. **(A)** Rescue of *ptc>dRet*^*M955T*^ fly lethality by ***1*** and *1-1.* Both showed improved efficacy (synergy) when co-administrated with 200 μΜ sorafenib (soraf). (−), vehicle DMSO control. Error bars represent standard error in triplicate experiments. *P < 0.05 in one-sided Student’s t-test as compared with vehicle control. **(B)** Docking pose of ***1*** and its analogs ***1-1*** and ***1-2*** (salmon sticks) with a DFG-out model of RET (broken yellow lines indicate hydrogen bonds), and their inhibition of migration of the dRet^M955T^-expressing cells. Right, suppression of cell migration by ***1*** and ***1-1***. Controls are shown in (C). **(C)** *In vivo* cell migration assay in *ptc>dRet*^*M955T*^ flies. Left, a developing whole wing disc containing GFP-labeled, dRet^M955T^-expressing cells constituting a stripe in the midline. The disc margin is visualized with DAPI (red pseudocolor). There are wild-type cells in black areas. Center, overgrowth of dRet^M955T^-expressing cells resulting in the thickening of the stripe in the apical view (top). In the z-series view (bottom), dRet^M955T^-expressing cells are migrating away from the original domain (arrows). Right, sorafenib suppressed the migration.

**Figure 5.**
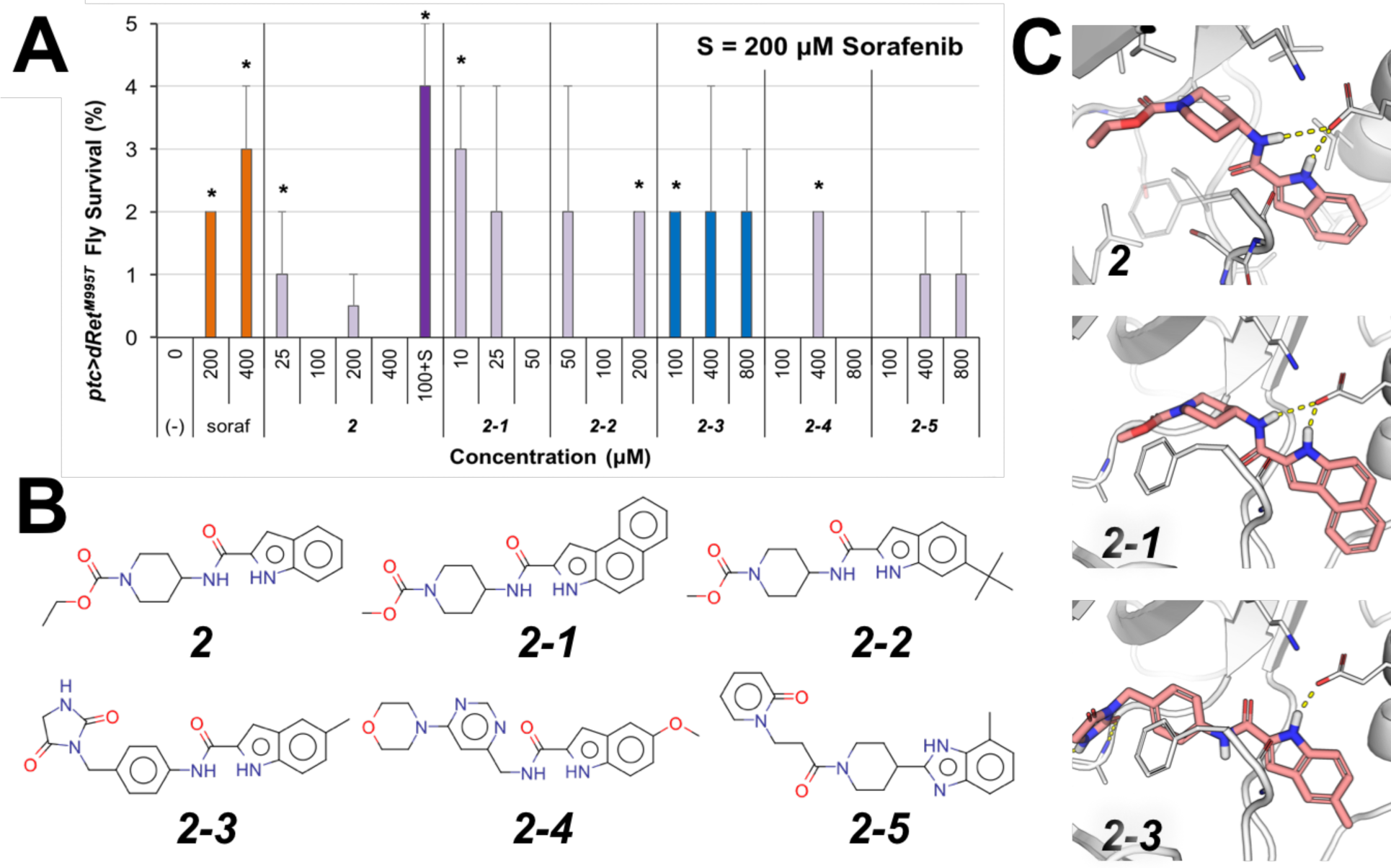
Rescue of *ptOdRef*^*9557*^ flies by 2 and its analogs. **(A)** viability assay. ***2*** showed increased efficacy when co-administrated with 200 μM sorafenib. (−), vehicle control. Error bars represent standard error in triplicate experiments. *P < 0.05 in one-sided Student’s t-test as compared with no-drug control. **(B)** Chemical structure of and ***2*** its analogs. **(C)** Docking pose of ***2*** and its analogs in a RET DFG-out model. These compounds are proposed to be putative type-II kinase inhibitors that bind in the DFG-pocket through their 1H-indole moiety and interact with the conserved aC-helix glutamate side chain and DFG-Aspartate backbone (broken yellow lines).

### Confirmation of novel chemical scaffolds

To validate the chemical scaffolds identified in our initial *Drosophila-based* chemical genetic screening, we conducted a ligand-based chemical similarity search in the updated ZINC [8] to identify analogs of ***1*** and **2**. For compound 1, we retrieved five compounds that share the 3-phenyl-(1H)-1,2,4-triazole feature and have docking poses similar to **1**. Our *ptc>dRet*^*M955T*^ screen confirmed two active compounds, ***1-1*** and ***1-2*** (Table S2; Fig. 4A, 4B). ***1-1*** outperformed *1* slightly in *ptc>dRet*^*M955T*^ fly rescue at similar concentrations (4%; *P* < 0.05). Conversely, ***1-2*** was tested at higher concentrations (50 and 200 μM) but did not result in better efficacy (P < 0.05).

The docking poses of ***1-1*** and ***1-2*** resemble the proposed docking pose of *1* (Fig. 4B), which has a typical DFG-out-specific, type-II KI binding pose and is predicted to occupy the DFG-pocket with the terminal phenyl moiety. The 1,2,4-triazole moiety, resembles the urea moiety found in sorafenib (Fig. 1A), forms favorable hydrogen bonds with the side chain of the conserved aC-helix glutamate residue and the backbone amide of the DFG-Aspartate. In addition, this series of compounds are smaller and shorter (MW < 360) than the fully developed type-II KIs (MW > 450) such as sorafenib, as they lack an optimized moiety that interacts with the hinge region of the ligand-binding site (Fig. 4C).

Compound ***1-2*** differs from ***1*** and ***1-1*** structurally and was less effective in rescuing *ptc>dRet*^*M955T*^ flies, even though it was tested at higher concentrations (Fig. 4A). While *1* and ***1-1*** have an (1H)-1,2,4-triazole moiety, ***1-2*** has an 1,2,4-oxadiazol-5-amine moiety, where the (1H)-nitrogen is replaced by an oxygen. This modification distinguishes ***1-2*** from ***1*** and ***1-1*** in their interaction preference: ***1-2*** loses a hydrogen bond donor due to the nitrogen-to-oxygen substitution, while the electronegative oxygen introduces an unfavorable electrostatic repulsion to the carboxylate sidechain of the conserved aC-helix glutamate (Fig. 4C, ***1-2*** insert).

Co-administering sorafenib with ***1*** and ***1-1*** led to synergistic improvement of *ptc>dRet*^*M955T*^ fly viability (Fig. 4A). Individually, 200 μM of sorafenib and 100 μM of ***1*** rescued 3% and 1% of *ptc>dRet*^*M955T*^ flies to adulthood, respectively. Co-administering the two compounds rescued 6% of *ptc>dRet*^*M955T*^ flies to adulthood (P < 0.05). Similarly, co-administering sorafenib and 100 μM of ***1-1*** rescued 8% of *ptc>dRet*^*M955T*^ flies (P < 0.05). In contrast, co-administering 200 μM of sorafenib and 200 μM of ***1-2*** did not improve fly viability. As ***1-2*** only weakly rescued *ptc>dRet*^*M955T*^ flies and showed no synergy with sorafenib, we did not pursue this hit any further.

We examined the kinase inhibition profile (DiscoverX) of ***1*** against a subset of the human protein kinome (Table 1). At 50 μM, *1* did not appreciably inhibit SRC, BRAF, or S6K1, while it demonstrated weak activity against wild-type RET and moderate activity against the oncogenic isoform RET^M918T^. Of note, *1* inhibited other cancer-related targets such as FLT3 (Table 1), which activates the Ras/ERK signaling pathway [22].

**Table 1.** Kinase inhibition profile of compound ***1*** at 50 μM.

| Kinase | % Inhib. | Kinase | % Inhib. |
| --- | --- | --- | --- |
| ABL1 | 0 | mTOR | 4 |
| AURKA | 22 | PDGFRB | 21 |
| AURKB | 7 | RET | 15 |
| AURKC | 2 | RET (M918T) | 28 |
| BRAF | 9 | RET (V804L) | 35 |
| CSF1R | 3 | S6K1 | 0 |
| FGFR | 0 | SRC | 5 |
| **FLT3** | **52** | TTK | 17 |

Bold, inhibited by more than 40%.

***1*** also showed activity against aspects of transformation and metastasis in the fly. In the mature larva, the *ptc* promoter is active in epithelial cells in a stripe pattern in the midline of the developing wing epithelium (Fig. 4C; wing disc). ptc-driven dRet activates multiple signaling pathways, promoting proliferation, epithelial-to-mesenchymal transition (EMT), and invasion of dRet^M955T^-expressing cells beyond the *ptc* domain [14] (Fig. 4C). Similar to sorafenib, oral administration of ***1*** blocked the invasion of dRet^M955T^-expressing cells into the surrounding wing epithelium (Fig. 4B).

At lower dosage (25 μM), compound ***2*** weakly rescued *ptc>dRet*^*M955T*^ flies (1%; *P* < 0.05) (Fig. 5A). Unlike **1, *2*** did not act synergistically with sorafenib. This difference was confirmed by the kinase inhibition profile of ***2*** (Table 2), in which it has stronger inhibition of RET and RET^M918T^, but loses the inhibition of FLT3, two key differences between the kinase inhibition profiles of ***1*** and **2**.

Through a chemical similarity search of the ZINC database, we identified five compounds that share the 1H-indole-2-carboxamide moiety with docking poses similar to that of ***2*** (Fig. 5B, 5C; Table S2), and confirmed all five analogs were able to improve the viability of *ptc>dRet*^*M955T*^ flies (Fig. 5A), albeit with weak efficacy (some have P-value above 0.05). At low dose (10 μM), ***2-1*** showed improved efficacy in rescuing *ptc>dRet*^*M955T*^ flies relative to ***2*** and had similar efficacy as sorafenib at 200 μM. However, ***2-1*** showed poor solubility, limiting its usefulness as lead compound. ***2-3*** was also more efficacious than *2* and displayed better solubility in both DMSO and water than ***2-1;*** it also has the W-phenylacetamide moiety as a linker group, a common linker feature found in type-II KIs such as imatinib. Compound ***2-3*** displayed a different inhibition profile than ***1*** and ***2*** (Table 2): it strongly inhibits FLT3 and PDGFRB, though is weak against both RET and ret^M918T^ and does not inhibit SRC.

**Table 2.** Kinase inhibition profile of compounds ***2*** and ***2-3*** at 50 μM.

| Compound **2** |  |  |  | Compound **2-3** |  |  |  |
| --- | --- | --- | --- | --- | --- | --- | --- |
| Kinase | % Inhib. | Kinase | % Inhib. | Kinase | % Inhib. | Kinase | % Inhib. |
| ABL1 | 0 | mTOR | 5 | ABL1 | 2 | mTOR | 8 |
| AURKA | 10 | PDGFRB | 20 | AURKA | 13 | PDGFRB | **91** |
| AURKB | 2 | RET | 34 | AURKB | 22 | RET | 24 |
| AURKC | 22 | **RET (M918T)** | **44** | AURKC | 2 | RET (M918T) | 23 |
| BRAF | 0 | **RET (V804L)** | **44** | BRAF | 0 | **RET (V804L)** | **43** |
| CSF1R | 12 | S6K1 | 0 | CSF1R | 20 | S6K1 | 0 |
| FGFR | 0 | SRC | 6 | FGFR | 4 | SRC | 0 |
| FLT3 | 25 | TTK | 31 | **FLT3** | **78** | TTK | 26 |

Bold, inhibited by more than 40%.

### Improving efficacy through compound hybridization

Interestingly, the chemical scaffolds of our newly identified active compounds are not associated with inhibition of protein kinases, as the analysis with SEA search [23] — which relates ligand chemical similarity of ligands to protein pharmacology — suggests. However, they provide rescue of *ptc>dRet*^*M955T*^ flies similar to that of sorafenib and regorafenib [15]. The docking poses of these compounds suggest a less-than-optimal interaction with the hinge region of protein kinases, a common feature of most KIs. Furthermore, the relatively low molecular weight (~350 g/mol) of these lead-like compounds provides a window for conducting lead optimization with medicinal chemistry. Hence, we sought to improve the efficacy of our computationally derived leads by installing a hinge-binding moiety found in known type-II KIs such as sorafenib.

We took into consideration the docking poses and phenotypic results of the known type-II kinase inhibitors (sorafenib and AD80 [1]) and lead compounds, as well as the synthetic accessibility and the novelty of the putative hybrid compounds, even if they do not dock optimally to our intended kinase targets. We focused on the functionalization of ***2/2-3*** based on these observations: 1) their 1H-indole moiety docks uniquely into the DFG-pocket and with the potential to interact with the aC-helix glutamate (Fig. 5C), 2) their 1H-indole-2-carboxamide moiety resembles the urea linker that is commonly found in type-II KIs such as sorafenib (Fig. 6A; blue box), and 3) the W-phenylcarboxamide moiety of ***2-3*** is a common linker between the hinge-binding and the DFG-pocket moieties of type-II KIs, e.g. imatinib (Fig. 6A; grey box), while the W-(piperidin-4-yl)carboxamide moiety of ***2*** is not a common linker, and **4**) docking pose of **2/3**’s 1H-indole moiety overlaps with the trifluoromethylphenyl moiety of sorafenib/AD80. We performed a fragment exchange at the carboxamide position by combining the 1H-indole-2-carboxamide moiety of ***2/2-3*** with the hinge-binding moiety of sorafenib and of AD80, a multi-kinase inhibitor that has shown promise in MTC treatment [1], to create ***3*** and ***4***, respectively (Fig. 6B).

**Figure 6.**
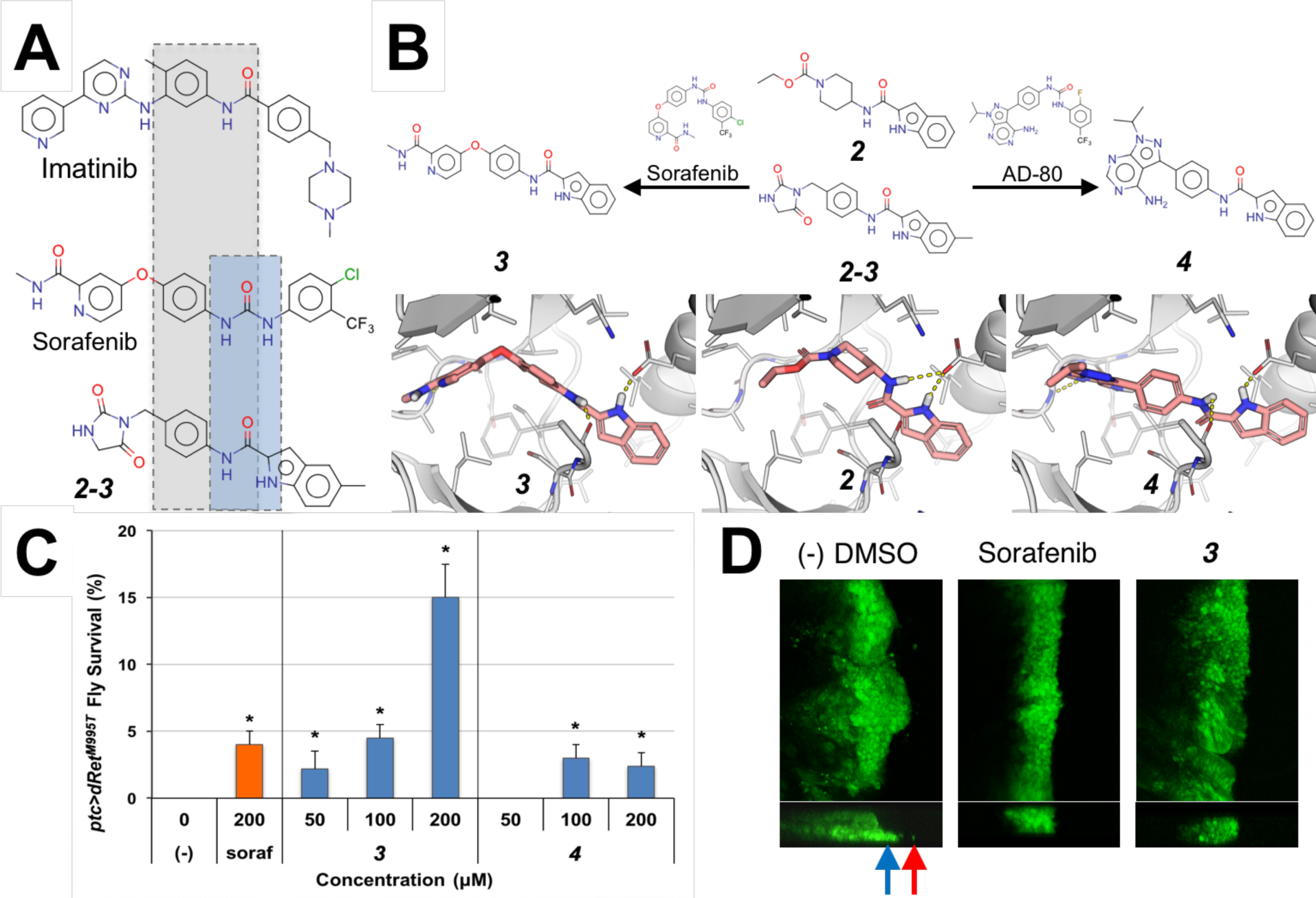
Hybrid compounds with improved efficacy. **(A)** The kinase inhibitors imatinib, sorafenib, and ***2-3*** share the common W-phenylcarboxamide moiety (grey box), while the 1H-indole-2-carboxamide of ***2-3*** resembles the urea linker of sorafenib (blue box). **(B)** Hybridization of ***2*** and sorafenib and AD-80 yielded ***3*** and ***4***, respectively. Top, chemical structures of compounds. Bottom, docking poses of compounds in a RET DFG-out model. **(C) *3*** rescued *ptc>dRet*^*M955T*^ flies more effectively than by either ***2*** or sorafenib alone. (−), vehicle control. Error bars represent standard error in triplicate experiments. *P > 0.05 in one-sided Student’s t-test as compared with no-drug control. **(D) *3*** suppresses migration of dRet^M955T^-expressing wing disc cells from the original domain (green) similarly to the positive control, sorafenib. Top and bottom, apical and z-series views, respectively. Arrows, migrating cells.

Oral administration of ***3*** and ***4*** to *ptc>dRet*^*M955T*^ flies demonstrated that the efficacy of ***4*** was low with only 3% rescue, while *3* demonstrated much improved efficacy with 15% rescue (Fig. 6C; *P* < 0.05), significantly higher rescue than the parent compound ***2/2-3*** and sorafenib. Additionally, ***3*** suppressed the invasion/migration of dRet^M955T^-expressing cells in the wing epithelium (Fig. 6D), further confirming its efficacy against dRet^M955T^-induced oncogenicity. The kinase inhibition profile of ***3*** (Table 3) resembles that of the parent compound ***2-3*** (Table 2) with at least two notable exceptions: ***3*** inhibits CSF1R, PDGFRB, and FLT3, all are receptor tyrosine kinases and orthologs of *Drosophila* Pvr that activate the Ras/ERK signaling pathway [24] and play key roles in SRC activation and tumor progression; and the inhibition of Aurora kinases AURKB and AURKC *(Drosophila* ortholog aurA or aurB). Of note, although ***4*** did not improve the viability of *ptc>dRet*^*M955T*^ flies, it shares chemical similarity to several known type-11 /2 kinase inhibitors that have the common adenine moiety and a related indole moiety. This group of inhibitors was shown to inhibit other related kinases, increasing our confidence in the relevance of this chemical space for kinase pathway modulation [25].

**Table 3.** Kinase inhibition profile of compound ***3*** at 50 μM.

| Kinase | % Inhib. | Kinase | % Inhib. |
| --- | --- | --- | --- |
| ABL1 | 0 | mTOR | 4 |
| AURKA | 15 | **PDGFRB** | **98** |
| **AURKB** | **88** | RET | 29 |
| **AURKC** | **91** | RET (M918T) | 26 |
| BRAF | 4 | **RET (V804L)** | **46** |
| **CSF1R** | **95** | S6K1 | 0 |
| FGFR | 0 | SRC | 4 |
| **FLT3** | **80** | TTK | 27 |

Bold, inhibited by more than 40%.

## DISCUSSION

### Integrated discovery pipeline

This study demonstrates the utility of an integrated platform that combines *Drosophila* genetics, computational structural biology, and chemical synthesis to enrich for the discovery of useful chemical tools in an established *Drosophila* MTC model (Fig. 2). We have previously shown that *Drosophila* can provide a unique entry point for drug development, by capturing subtle structural changes in lead compounds that are often missed by cellular or biochemical assays. Here we refine this approach by iteratively combining experimental testing with computational modeling. Overall, a key strength of the combined approach is its ability to rapidly and in a cost-effective manner test chemically unique, purchasable compounds with our fly models; this platform allowed us to quickly confirm the *in situ* relevance of active chemotypes through iterations of computational modeling, synthetic chemistry, and phenotypic testing in the fly. We expect this integrated pipeline is generally applicable to kinase networks associated with other diseases [7].

### DFG-out modeling approach

DFGmodel is a recent computational development that generates models of kinases in their inactive, DFG-out conformation for rational design of type-II KIs [10]. In a recent study, models generated by DFGmodel were used to guide the optimization of the drug sorafenib, to target a new disease space [15]. Here, we demonstrated a successful application of DFGmodel to explore compounds that are not appreciated as kinase inhibitors. For each kinase target, DFGmodel uses multiple experimentally determined structures as modeling templates and generates multiple homology models. Thus, this method samples a large fraction of the DFG-out conformational space during the model construction, which enables us to account for the flexibility of the binding site during virtual screening [26]. Notably, DFG-out models capture key features that are important for protein-ligand interactions in multiple kinases simultaneously, providing a framework for rationalizing activity of known inhibitors and developing unique KIs that target a signaling pathway. For example, our results suggest that the electrostatic potential within the DFG-pocket is a key feature for inhibitor selectivity: ERK has an inverse electrostatic potential in the DFG-pocket than that of the target kinases such as RET and BRAF (Fig. 3), which may explain the insensitivity of ERK toward inhibitors such as sorafenib.

### Identification of biologically active compounds

Most clinically approved KIs are ineffective against MTC; the most effective inhibitors, sorafenib and regorafenib, show limited efficacy in the *ptc>dRet*^*M955T*^ fly model, rescuing 3-4% at 200 μM [15]. Despite considerable academic and industry effort, the known chemical space of kinase inhibitors is limited [7]. For example, sorafenib and regorafenib differ in only one non-hydrogen atom. Through structure-based virtual screening against multiple kinase targets in a disease pathway, we discovered chemically unique compounds (Table S2) with an ability to rescue *ptc>dRet*^*M955T*^ viability that is similar to the most effective FDA-approved drug sorafenib (Figs. 4 and 5).

Importantly, our data indicates that these compounds act through key cancer networks. For example, compounds ***1, 2, 2-3*** and ***3*** all have shown the ability to suppress invasion of *ptc>dRet*^*M955T*^*cells* in the wing epithelium. Previous works demonstrated that wing cell invasion is controlled by SRC [15, 27], which acts by controlling E-cadherin and Matrix Metalloproteases (MMPs). Of note, ***1, 2, 2-3*** and ***3*** each show significant activity against orthologs of *Drosophila* Pvr, a key regulator of Src: all show significant activity against human FLT3, while *3* shows additional activity against Pvr orthologs CSF1R and PDGFRB. In addition to being orthologs of Pvr, FLT3, CSF1R, and PDGFRB similarly can activate SRC [28]. We propose that this activity against regulators of SRC account for the ability of ***1, 2, 2-3*** and ***3*** to suppress invasion, a key first step in tumor metastasis. Other activities, for example, ***3***’s inhibition of Aurora kinases — required for proliferation during tumor progression [29] — likely also contributes. Indeed, AURK inhibitors are known to be active against MTC [30, 31] and synergy between AURKs and FLT3 is currently being explored clinically through a number of dual-AURKB/FLT3 inhibitors [32] [33].

### Recombination of building blocks for future inhibitors

Though the new tool compounds ***1*** and ***2*** may not be sufficiently potent to serve as therapeutics, they reveal diverse fragment-like pharmacophores that serve as starting points for an exploration of new chemical space. These pharmacophores can be further optimized by combining with well-developed chemotypes that are known to interact with kinase binding sites (e.g., the hinge binding region) to form more efficacious chemical probes; this provides a key second step towards building effective compounds. For example, ***2*** and ***2-3*** include an 1H-indole moiety capable of occupying the DFG-pocket of protein kinases from different families and a carboxamide group commonly found in type-II KIs (Fig. 6A). Guided by the docking poses of these compounds, the 1H-indole-2-carboxamide group was combined with an optimized hinge-binding moiety from sorafenib, to form a significantly more efficacious compound (i.e., **3**). As indicated in the kinase inhibition profile of ***3*** (Table 3), it shares part of the target set of its constituents ***2*** and ***2-3***.

In summary, we demonstrate the potential of combining chemical modeling with *Drosophila* genetics to rapidly and efficiently explore novel chemical space. This provides an accessible and cost-effective platform that can be applied to a broad palette of diseases that can be modeled in *Drosophila.* Combining the strengths of these two high-throughput approaches opens the opportunity to develop novel tool and lead compounds that are effective in the context of the whole animal.

## MATERIALS & METHODS

### DFG-out models

Models of kinase targets (human RET, SRC, BRAF, S6K1) in the DFG-out conformation were generated using DFGmodel [10]. Briefly, the method takes a DFG-in structure or the sequence of the protein kinase as input. DFG-model relies on a manually curated alignment between the target kinase and multiple structures representing unique DFG-out conformations. It calls on the structure-based sequence alignment function of T_COFFEE/Expresso [34] v11.00.8 to perform sequence alignment of the kinase catalytic domain to the templates, followed by the multi-template function of MODELLER [35] v9.14 to generate 50 homology models covering a range of conformations. For each kinase 10 DFG-out models with largest binding site volume, as calculated by POVME [36] v2.0, were used for further study. These ensembles of models were evaluated and confirmed to enrich known type-II inhibitors over non-ligands using docking, which provides an approximation of the binding site flexibility, as well as optimizes the binding site for protein-ligand complementarity and structure-based virtual screening [11, 26, 37]. The area-under-curve (AUC) of targets BRAF and RET DFG-out models are 90.6 vs. 82.8, respectively, which correspond to at least 5-fold increase in the enrichment accuracy over randomly selected ligands in a known sample set [37–39].

### Virtual screening

Initial virtual screening utilized the ZINC12 [21] “available now” leadlike chemical library (downloaded in 2013, 2.2 million compounds). Default settings were used for the ligand conformer generation with OMEGA and the docking program FRED [40][41]. For each kinase, the ensemble of 10 DFG-out models was used for screening and the results were processed with the open-source cheminformatics toolkit RDKit (www.rdkit.org). To obtain a consensus docking result for RET and the targets BRAF, SRC, p70-S6K, a two-step approach was used: 1) 2,000 ligands were collected for each kinase by identifying ligands that ranked in the top 10% for at least half of the DFG-out ensemble.; 2) ligands that scored in the top 25% in at least 3 of 4 kinase ensembles were collated into a final set of 247 compounds. These consensus ligands, representing 0.0114% of the library, were visually inspected to remove molecules with energetically unfavorable or strained conformations, or with reactive functional groups that may interfere with assays [41], which are commonly observed in large virtual screenings. 8 compounds were selected based on their interactions with the receptor (DFG-pocket occupancy, hydrogen-bond to conserved amino acids, etc) and chemical novelty and were purchased for *Drosophila* viability screening. Analogs ***1-1, 1-2, 2-1 to 2-5***, and others (Table S1) were identified based on the structure of compounds *1* and *2* through the chemical similarity search function available in ZINC15 [8] and SciFinder using the default setting and Tanimoto coefficient above 70%. These compounds are commercially available through vendors such as ChemBridge and Enamine.

### Chemical Methods

For synthetic procedures and characterization data related to compounds ***1, 3***, and ***4***, please see supplementary materials.

### Kinase profiling of compounds

Kinase inhibition profile of the compounds was assessed at 50 μM through commercially available kinase profiling services (DiscoverX).

### Drosophila stocks

Human orthologs of *Drosophila* genes were predicted by DIOPT (http://www.flyrnai.org/cgi-bin/DRSC_orthologs.pl). The multiple endocrine neoplasia (MEN) type 2B mutant form of *Drosophila* Ret carries the M955T mutation (dRet^M955T^), which corresponds to the M918T mutation found in human MTC patients. The *ptc-gal4, UAS-GFP; UAS-dRet*^*M955T*^*/SM5(tub-gal80)-TM6B* transgenic flies were prepared according to standard protocols [15]. In these flies, the *tubulin* promoter drives GAL80, a suppressor of GAL4, to repress dRet^M955T^ expression. We crossed them with *w*^−^ flies to obtain *ptc>dRet*^*M955T*^flies that lost *GAL80* allele, which derepressed dRet^M955T^ expression (Fig. S1A). Transgenic *ptc>Ret*^*M955T*^ flies were calibrated to have 0% survival when raised at 25°C, which allowed for drug screening (Fig. S1B).

### Chemical genetic screening in flies

We employed dominant modifier screening [15] using the *ptc-gal4, UAS-GFP; UAS-dRet*^*M955T*^*/SM5(tub-gal80)-TM6B* to screen for fly kinase genes that affected the *dRet*^*M955T*^*-induced* lethality in flies when heterozygous *(ptc>Ret^M955T^;kinase^+^).* Genes that improved or reduced survival of *ptc>dRet*^*M955T*^ flies when heterozygous were designated as genetic ‘suppressors’ or ‘enhancers’, respectively (Fig. 1B). Suppressors are candidate therapeutic targets that when inhibited may reduce tumor progression.

Stock solutions of the test compounds were created by dissolving the compound in DMSO at the maximum concentration. The stock solutions were diluted by 1000-fold or more and mixed with semi-defined fly medium (Bloomington Drosophila Stock Center) to make drug-infused food (0.1% final DMSO concentration; maximum tolerable dose in flies). Approximately 100 *ptc>dRet*^*M955T*^ embryos alongside with wild-type *(+;+/SM5tubgaleo-TM6B)* flies were raised until adulthood on drug-infused food for 13 days at 25°C. The numbers of empty pupal cases (P in Fig. S1B) and that of surviving adults (A) were used to determine percentage of viability, while their body size, which is affected by food intake, temperature, and humidity, were compared to vehicle-treated groups to standardize the experimental conditions.

### Wing discs cell migration/invasion assays

Third-instar *ptc>dRet*^*M955T*^ larvae were dissected, and developing wing discs were collected, fixed with 4% paraformaldehyde in PBS, and whole-mounted. At least 10 wing discs were analyzed for each treatment. Invasive GFP-labeled dRet^M955T^-expressing cells were visualized by their green pseudocolor under a confocal microscope. The apical and the virtual z-series views of the wing disc were examined to identify abnormal tissue growth and dRet^M955T^-expressing cells migrating beyond the *ptc* domain boundary.

## ACKNOWLEDGMENTS

We thank Peter Smibert (New York Genome Center) for *ptc>dRet*^*M955T*^ flies, Kevin Cook (Bloomington *Drosophila* Stock Center) for kinome mutant fly lines. This work was supported by the National Institutes of Health (R01-GM108911) to A.S. and P.M.U.U. A.S. was also supported by Department of Defense grant W81XWH-15-1-0539. M.S. and R.C. were supported by National Institutes of Health grants U54OD020353, R01-CA170495, and R01-CA109730 and Department of Defense grant W81XWH-15-1-0111. The Dar laboratory (A.P.S. and A.C.D.) is supported by Innovation awards from the NIH (DP2 CA186570-01) and Damon Runyon-Rachleff Foundation. A.C.D. is a Pew-Stewart Scholar in Cancer Research and Young Investigator of the Pershing-Square Sohn Cancer Research Alliance. We appreciate OpenEye Scientific Software, Inc. for granting us access to its high-performance molecular modeling applications through its academic license program. This work was supported in part through the computational resources and staff expertise provided by the Department of Scientific Computing at the Icahn School of Medicine at Mount Sinai.

## AUTHOR CONTRIBUTIONS

P.M.U.U. performed and analyzed the homology modeling of kinases, virtual screening of compound library, selection and design of testing compounds. M.S. managed *Drosophila* stocks and testing compounds in whole animal and *in vivo* experiments. A.P.S. conducted organic synthesis, design, and validation of test compounds. P.M.U.U. analyzed results and wrote the manuscript with input from all co-authors. R.L.C., A.C.D., and A.S. initiated, supervised, and acquired funding and resources for the project.

## SUPPORTING INFORMATION LEGENDS

**Figure S1. (A)** Preparation of transgenic *ptc>dRet*^*M955T*^ flies for chemical genetic screening [3]. **(B)** Determination of compound efficacy in a fly-based chemical genetic screening. The numbers of empty pupal cases (P) and surviving adult *(A)* are used to determine viability.

**Figure S2. DFG-pocket of various protein kinases.** The left panels show the DFG-pocket (colored volume) with the docking pose of ***1.*** The right panels show the electrostatic potential on the surface of DFG-pocket (blue, positive; red, negative).

**Figure S3. Common interactions in type-II inhibitor binding site.** Type-II kinase inhibitors are modular. They are composed of a hinge-binding moiety and a spacer group, followed by a linker that forms hydrogen bonds with the conserved glutamate residue on the aC-helix, as well as a hydrophobic “cap” group that docks into the DFG-pocket. Key elements in type-II inhibitor/kinase interactions include **(A)** Hydrogen bonds with “hinge” amide backbone. **(B)** π-π stacking with DFG-Phe. **(C)** Hydrogen bonds with aC-helix glutamate. **(D)** Hydrogen bond with DFG-Asp amide backbone. **(E)** van der Waals interactions in DFG-pocket.

## REFERENCES

1. Dar AC, Das TK, Shokat KM, Cagan RL. Chemical genetic discovery of targets and anti-targets for cancer polypharmacology. Nature. 2012;486(7401):80–4. doi: 10.1038/nature11127.

2. Kasai Y, Cagan R. Drosophila as a tool for personalized medicine: a primer. Personalized medicine. 2010;7(6):621–32. doi: 10.2217/pme.10.65.

3. Sonoshita M, Cagan RL. Modeling Human Cancers in Drosophila. Curr Top Dev Biol. 2017;121:287–309. doi: 10.1016/bs.ctdb.2016.07.008.

4. Vidal M, Wells S, Ryan A, Cagan R. ZD6474 suppresses oncogenic RET isoforms in a Drosophila model for type 2 multiple endocrine neoplasia syndromes and papillary thyroid carcinoma. Cancer research. 2005;65(9):3538–41. doi: 10.1158/0008-5472.CAN-04-4561.

5. Manning G, Whyte DB, Martinez R, Hunter T, Sudarsanam S. The protein kinase complement of the human genome. Science. 2002;298(5600):1912–34. doi: 10.1126/science.1075762.

6. Read RD, Goodfellow PJ, Mardis ER, Novak N, Armstrong JR, Cagan RL. A Drosophila model of multiple endocrine neoplasia type 2. Genetics. 2005;171(3):1057–81. doi: 10.1534/genetics.104.038018.

7. Schlessinger A, Abagyan R, Carlson HA, Dang KK, Guinney J, Cagan RL. Multitargeting Drug Community Challenge. Cell Chem Biol. 2017;24(12):1434–5. doi: 10.1016/j.chembiol.2017.12.006.

8. Sterling T, Irwin JJ. ZINC 15—Ligand Discovery for Everyone. Journal of chemical information and modeling. 2015;55(11):2324–37. doi: 10.1021/acs.jcim.5b00559.

9. Irwin JJ, Shoichet BK. Docking Screens for Novel Ligands Conferring New Biology. Journal of medicinal chemistry. 2016;59(9):4103–20. doi: 10.1021/acs.jmedchem.5b02008.

10. Ung PMU, Schlessinger A. DFGmodel: predicting protein kinase structures in inactive states for structure-based discovery of type-II inhibitors. ACS chemical biology. 2015;10(1):269–78. doi: 10.1021/cb500696t.

11. Kufareva I, Abagyan R. Type-II kinase inhibitor docking, screening, and profiling using modified structures of active kinase states. Journal of medicinal chemistry. 2008;51(24):7921–32. doi: 10.1021/jm8010299.

12. Cerrato A, De Falco V, Santoro M. Molecular genetics of medullary thyroid carcinoma: the quest for novel therapeutic targets. Journal of molecular endocrinology. 2009;43(4):143–55. doi: 10.1677/JME-09-0024.

13. Hadoux J, Pacini F, Tuttle RM, Schlumberger M. Management of advanced medullary thyroid cancer. The lancet Diabetes & endocrinology. 2016;4(1):64–71. doi: 10.1016/S2213-8587(15)00337-X.

14. Vidal M, Larson DE, Cagan RL. Csk-deficient boundary cells are eliminated from normal Drosophila epithelia by exclusion, migration, and apoptosis. Developmental cell. 2006;10(1):33–44. doi: 10.1016/j.devcel.2005.11.007.

15. Sonoshita M, Scopton AP, Ung PMU, Murray MA, Silber L, Maldonado AY, et al. A whole-animal platform to advance a clinical kinase inhibitor into new disease space. Nature chemical biology. 2018;14(3):291–8. doi: 10.1038/nchembio.2556.

16. Wan PT, Garnett MJ, Roe SM, Lee S, Niculescu-Duvaz D, Good VM, et al. Mechanism of activation of the RAF-ERK signaling pathway by oncogenic mutations of B-RAF. Cell. 2004;116(6):855–67. doi: 10.1016/S0092-8674(04)00215-6.

17. Seeliger MA, Nagar B, Frank F, Cao X, Henderson MN, Kuriyan J. c-Src binds to the cancer drug imatinib with an inactive Abl/c-Kit conformation and a distributed thermodynamic penalty. Structure. 2007;15(3):299–311. doi: 10.1016/j.str.2007.01.015.

18. Huse M, Kuriyan J. The conformational plasticity of protein kinases. Cell. 2002;109(3):275–82. doi: 10.1016/S0092-8674(02)00741-9.

19. Zhao Z, Wu H, Wang L, Liu Y, Knapp S, Liu Q, et al. Exploration of type II binding mode: A privileged approach for kinase inhibitor focused drug discovery? aCs chemical biology. 2014;9(6):1230–41. doi: 10.1021/cb500129t.

20. Mulligan LM. RET revisited: expanding the oncogenic portfolio. Nature reviews Cancer. 2014;14(3):173–86. doi: 10.1038/nrc3680.

21. Irwin JJ, Sterling T, Mysinger MM, Bolstad ES, Coleman RG. ZINC: a free tool to discover chemistry for biology. Journal of chemical information and modeling. 2012;52(7):1757–68. doi: 10.1021/ci3001277.

22. Scholl C, Gilliland DG, Frohling S. Deregulation of signaling pathways in acute myeloid leukemia. Semin Oncol. 2008;35(4):336–45. doi: 10.1053/j.seminoncol.2008.04.004.

23. Keiser MJ, Roth BL, Armbruster BN, Ernsberger P, Irwin JJ, Shoichet BK. Relating protein pharmacology by ligand chemistry. Nat Biotechnol. 2007;25(2):197–206. doi: 10.1038/nbt1284.

24. Schlessinger J. Cell signaling by receptor tyrosine kinases. Cell. 2000;103(2):211–25 doi: 10.1016/S0092-8674(00)00114-8.

25. Burchat A, Borhani DW, Calderwood DJ, Hirst GC, Li B, Stachlewitz RF. Discovery of A-770041, a src-family selective orally active lck inhibitor that prevents organ allograft rejection. Bioorganic & medicinal chemistry letters. 2006;16(1):118–22. doi: 10.1016/j.bmcl.2005.09.039.

26. Amaro RE, Li WW. Emerging methods for ensemble-based virtual screening. Curr Top Med Chem. 2010;10(1):3–13. doi: 10.2174/156802610790232279.

27. Vidal M, Warner S, Read R, Cagan RL. Differing Src signaling levels have distinct outcomes in Drosophila. Cancer research. 2007;67(21):10278–85. doi: 10.1158/0008-5472.CAN-07-1376.

28. Sachsenmaier C, Sadowski HB, Cooper JA. STAT activation by the PDGF receptor requires juxtamembrane phosphorylation sites but not Src tyrosine kinase activation. Oncogene. 1999;18(24):3583–92. doi: 10.1038/sj.onc.1202694.

29. Sasai K, Katayama H, Stenoien DL, Fujii S, Honda R, Kimura M, et al. Aurora-C kinase is a novel chromosomal passenger protein that can complement Aurora-B kinase function in mitotic cells. Cell Motil Cytoskeleton. 2004;59(4):249–63. doi: 10.1002/cm.20039.

30. Baldini E, Arlot-Bonnemains Y, Sorrenti S, Mian C, Pelizzo MR, De Antoni E, et al. Aurora kinases are expressed in medullary thyroid carcinoma (MTC) and their inhibition suppresses in vitro growth and tumorigenicity of the MTC derived cell line TT. BMC Cancer. 2011;11:411. doi: 10.1186/1471-2407-11-411.

31. Tuccilli C, Baldini E, Prinzi N, Morrone S, Sorrenti S, Filippini A, et al. Preclinical testing of selective Aurora kinase inhibitors on a medullary thyroid carcinoma-derived cell line. Endocrine. 2016;52(2):287–95. doi: 10.1007/s12020-015-0700-0.

32. Bavetsias V, Linardopoulos S. Aurora Kinase Inhibitors: Current Status and Outlook. Front Oncol. 2015;5:278. doi: 10.3389/fonc.2015.00278.

33. Grundy M, Seedhouse C, Shang S, Richardson J, Russell N, Pallis M. The FLT3 internal tandem duplication mutation is a secondary target of the aurora B kinase inhibitor AZD1152-HQPA in acute myelogenous leukemia cells. Mol Cancer Ther. 2010;9(3):661–72. doi: 10.1158/1535-7163.MCT-09-1144.

34. Notredame C, Higgins DG, Heringa J. T-Coffee: A novel method for fast and accurate multiple sequence alignment. Journal of molecular biology. 2000;302(1):205–17. doi: 10.1006/jmbi.2000.4042.

35. Sali A, Blundell TL. Comparative protein modelling by satisfaction of spatial restraints. Journal of molecular biology. 1993;234(3):779–815. doi: 10.1006/jmbi.1993.1626.

36. Durrant JD, Votapka L, Sorensen J, Amaro RE. POVME 2.0: An Enhanced Tool for Determining Pocket Shape and Volume Characteristics. Journal of chemical theory and computation. 2014;10(11):5047–56. doi: 10.1021/ct500381c.

37. Fan H, Irwin JJ, Webb BM, Klebe G, Shoichet BK, Sali A. Molecular docking screens using comparative models of proteins. Journal of chemical information and modeling. 2009;49(11):2512–27. doi: 10.1021/ci9003706.

38. Katritch V, Rueda M, Lam PC, Yeager M, Abagyan R. GPCR 3D homology models for ligand screening: lessons learned from blind predictions of adenosine A2a receptor complex. Proteins. 2010;78(1):197–211. doi: 10.1002/prot.22507.

39. Carlsson J, Coleman RG, Setola V, Irwin JJ, Fan H, Schlessinger A, et al. Ligand discovery from a dopamine D3 receptor homology model and crystal structure. Nature chemical biology. 2011;7(11):769–78. doi: 10.1038/nchembio.662.

40. OEDOCKING 3.2.0.2: OpenEye Scientific Software, Santa Fe, ME.

41. Baell JB, Holloway GA. New substructure filters for removal of pan assay interference compounds (PAINS) from screening libraries and for their exclusion in bioassays. Journal of medicinal chemistry. 2010;53(7):2719–40. doi: 10.1021/jm901137j.

